## Supporting Information for "The primary mechanism for highly potent inhibition of HIV-1 maturation by lenacapavir"

**S1 Table. Production of HIV-1 particles.**

| pNL4.3, µg | p24 ng/ml |
|---|---|
| 0.03125 | 14.2 ± 0.5 |
| 0.125 | 153.6 ± 6.5 |
| 0.5 | 3504.3 ± 83.8 |
| 2.0 | 4690.5 ± 647.4 |

**S2 Table. Data collection and refinement statistics.**

| | $NTD_{LEN}$ (PDB ID: 8V23) |
|---|---|
| **Wavelength** | 1.072156 |
| **Resolution range** | 45.8 - 2.0 (2.072 - 2.0) |
| **Space group** | $P\ 2_1\ 2_1\ 2_1$ |
| **Unit cell** | 32.218 44.127 91.608 90 90 90 |
| **Total reflections** | 57809 (5804) |
| **Unique reflections** | 9274 (896) |
| **Multiplicity** | 6.2 (6.5) |
| **Completeness (%)** | 99.37 (100.00) |
| **Mean I/sigma(I)** | 18.82 (8.99) |
| **Wilson B-factor** | 25.61 |
| **R-pim** | 0.03553 (0.09962) |
| **CC1/2** | 0.996 (0.986) |
| **CC*** | 0.999 (0.996) |
| **Reflections used in refinement** | 9267 (896) |
| **Reflections used for R-free** | 975 (103) |
| **R-work** | 0.2301 (0.2524) |
| **R-free** | 0.2859 (0.2973) |
| **CC(work)** | 0.891 (0.637) |
| **CC(free)** | 0.898 (0.617) |
| **Number of non-hydrogen atoms** | 1187 |
| **macromolecules** | 1133 |
| **ligands** | 0 |
| **solvent** | 54 |
| **Protein residues** | 146 |
| **RMS(bonds)** | 0.009 |
| **RMS(angles)** | 1.05 |
| **Ramachandran favored (%)** | 96.53 |
| **Ramachandran allowed (%)** | 1.39 |
| **Ramachandran outliers (%)** | 2.08 |
| **Rotamer outliers (%)** | 1.64 |
| **Clashscore** | 3.98 |
| **Average B-factor** | 39.86 |
| **macromolecules** | 39.90 |
| **solvent** | 39.01 |

Statistics for the highest-resolution shell are shown in parentheses.

### Supplementary Figure Legends

**S1 Fig. Antiviral activity of DRV during late steps of HIV-1 replication.** HIV-1 virions were produced by transfecting indicated amounts of the full-length, WT HIV-1NL4.3 plasmid in HEK293T cells (producer cells). Indicated concentrations of DRV or DMSO control were added to HEK293T cells, the excess DRV was removed by the Lenti-XTM concentrator, and the virions were used to infect HeLa TZM-bl cells (Target cells). After 48 h of infection, luciferase activity was measured to determine the EC50 values for DRV. The averaged data (+/−SD) from three independent experiments are shown.

**S2 Fig. Effects of LEN on intracellular Gag levels.** HEK293T cells were transfected with 2 μg full-length, WT HIV-$1_{NL4.3}$ plasmid. After 4 h, increasing concentrations of LEN were added to the virus producer cells. After 48 h, cells were collected, and immunoblotting analysis was performed using anti-HIV1 p55 + p24 + p17 antibody (ab63917). GAPDH was used for internal control. Lane 1: molecular weight markers; lane 2: 2025 nM LEN; lane 3: 675 nM LEN; lane 4: 225 nM LEN; lane 5: 75 nM LEN; lane 6: 25 nM LEN; lane 7: 8.3 nM LEN; lane 8: 2.8 nM LEN; lane 9: DMSO control. The immunoblot is representative of results observed in two independent experiments.

**S3 Fig. LEN does not affect Gag proteolytic processing during HIV-1 particle maturation.** (A) HEK293T cells were transfected with 0.125 μg full-length WT HIV-1 NL4.3 plasmid. After 6 h post transfection, the medium was removed and 1 nM LEN or DMSO control containing media were added to the virus producer cells. The supernatants containing viruses were collected at 0 h, 16 h, 24 h, 40 h, 48 h, ultracentrifuged, and analyzed by immunoblotting. (B) HEK293T cells were transfected with 2 μg full-length WT HIV-1 NL4.3 plasmid. After 30 h post transfection, the medium was removed and 50 nM LEN or DMSO containing media were added to the virus producer cells. The supernatants containing virions were collected at 0 h, 2 h, 4 h, 8 h, ultracentrifuged, and analyzed by immunoblotting. Representative images of at least three independent experiments are shown in A and B. The observed Gag proteolytic processing products including CA, matrix (MA) and nucleocapsid (NC) are indicated.

**S4 Fig. Binding of LEN to recombinant full-length Gag.** (A) Representative SPR sensorgrams of LEN binding to full-length Gag showing association (crescent curve) and dissociation (decrescent curve); $K_D$, $k_{on}$ and $k_{off}$ values are indicated. (B) The $K_D$ value is determined using the Hill fit for (A).

**S5 Fig. Proteolytic processing of recombinant Gag by HIV-1 protease.** (A) HIV-1 protease mediated cleavage of full-length, recombinant Gag (1 μM) was performed in the presence of 2 μM LEN (+) or DMSO control (-). Lane 1: input of full-length Gag. Lanes 2 – 11: reaction products. Lane 12: MW markers. (B) HIV-1 protease mediated cleavage of 1 μM Gag(Δp6) was performed in the presence of 2 μM LEN (+) or DMSO control (-). Lane 1: input of Gag(Δp6). Lanes 2 – 11: reaction products. Lane 12: MW markers. Representative SDS-PAGE images of at least three independent reactions are shown. The proteolytic products including the bands corresponding to p41, CA, MA and NC are indicated.

**S6 Fig. Comparison of modelled WT CA pentamer with the cross-linked CA pentamer.** Superimposition of the AlphaFold2 generated CA pentamer structure (orange) and X-ray crystallographic structure of the cross-linked pentamer (PDB id: 3PO5, in purple); (A) top and (B) side views.

**S7 Fig. Comparative analysis of the opened and closed conformations of CA NTD.** The distances between C$\alpha$ of G60 and Cε of M66 are shown to delineate the opened and closed conformations. (A) $NTD_{LEN}$ (PDB: 8V23, present work); (B) $CA_{Hex}$ (PDB: 7URN); (C) $CA_{Hex}$ + LEN (PDB 6VKV); (D) NTD + PF74 (PDB: 2XDE); (E) $NTD_{Apo}$ (PDB: 5HGK); (F) $CA_{Pnt}$ (PDB: 7URN); (G) NTD + CAP-1 (PDB: 2JPR); (H) NTD + 1F6 (PDB: 4INB). Colored protein segments indicate connecting helices H3 and H4. Black dash lines indicate distances between C$\alpha$ of Gly60 and Cε of Met66.

**S8 Fig. LEN binding is compatible with the opened but not the closed conformation of CA.** Structure of $CA_{Hex}$ (Green) + LEN (dark blue) (PDB: 6VKV) superimposed onto (A) NTD + LEN ($NTD_{LEN}$, steel blue, PDB: 8V23, present work); (B) $CA_{Pnt}$ (scarlet, PDB: 7URN); and (C) $NTD_{Apo}$ (magenta, PDB: 5HGK). Different conformations of Met66 side chains are shown. In panel A, the Met66 side chain is compatible with LEN binding to the opened conformation seen in $NTD_{LEN}$. In contrast, the Met66 side chains in native $CA_{Pnt}$ (B) and $NTD_{Apo}$ (C) encounters steric hindrance with respect to LEN.

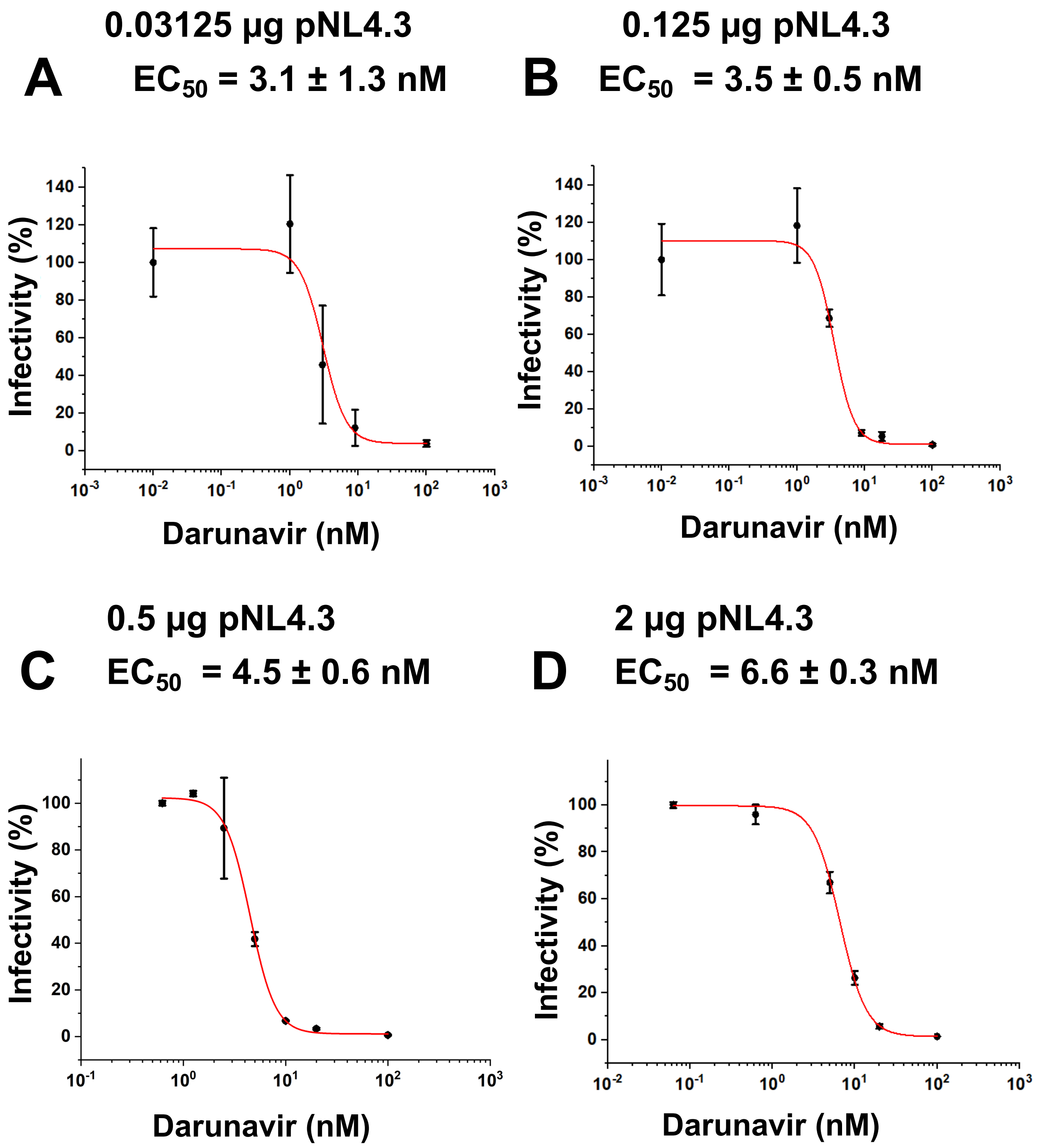
A
0.03125 µg pNL4.3
EC50 = 3.1 ± 1.3 nM
B
0.125 µg pNL4.3
EC50 = 3.5 ± 0.5 nM
C
0.5 µg pNL4.3
EC50 = 4.5 ± 0.6 nM
D
2 µg pNL4.3
EC50 = 6.6 ± 0.3 nM
Infectivity (%)
Darunavir (nM)

S1 Fig

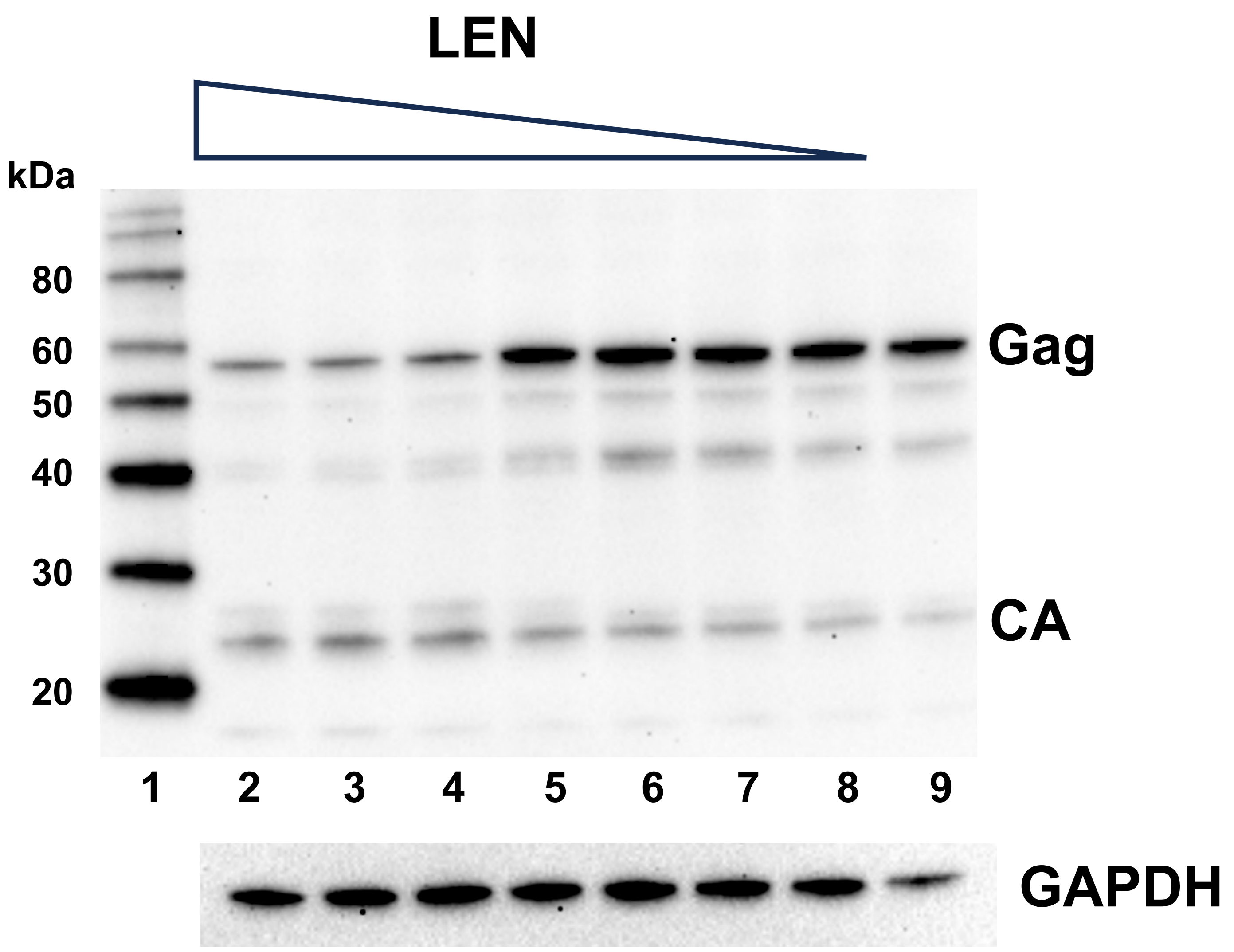


S2 Fig

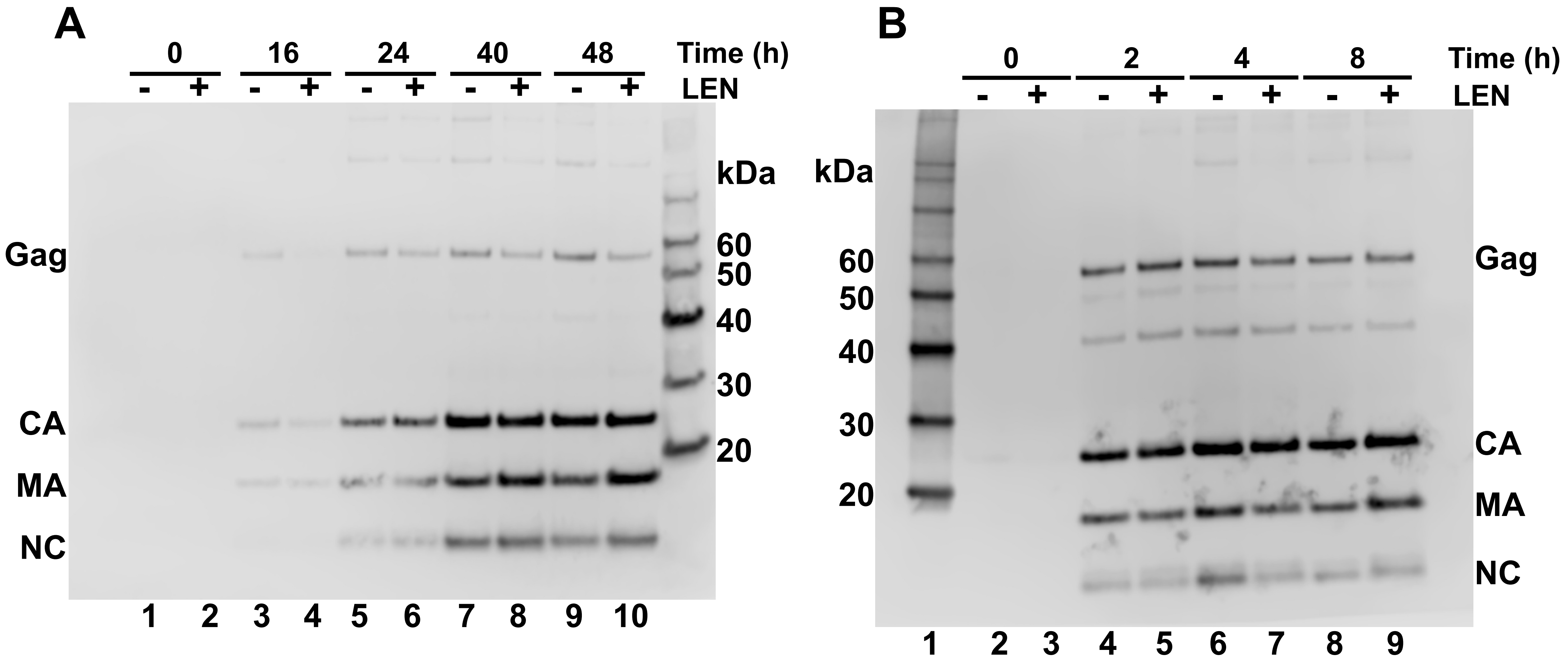
A
0
16
24
40
48
Time (h)
- + - + - + - + - +
LEN
Gag
CA
MA
NC
kDa
60
50
40
30
20
1 2 3 4 5 6 7 8 9 10
B
0
2
4
8
Time (h)
- + - + - + - +
LEN
kDa
60
50
40
30
20
Gag
CA
MA
NC
1 2 3 4 5 6 7 8 9

S3 Fig

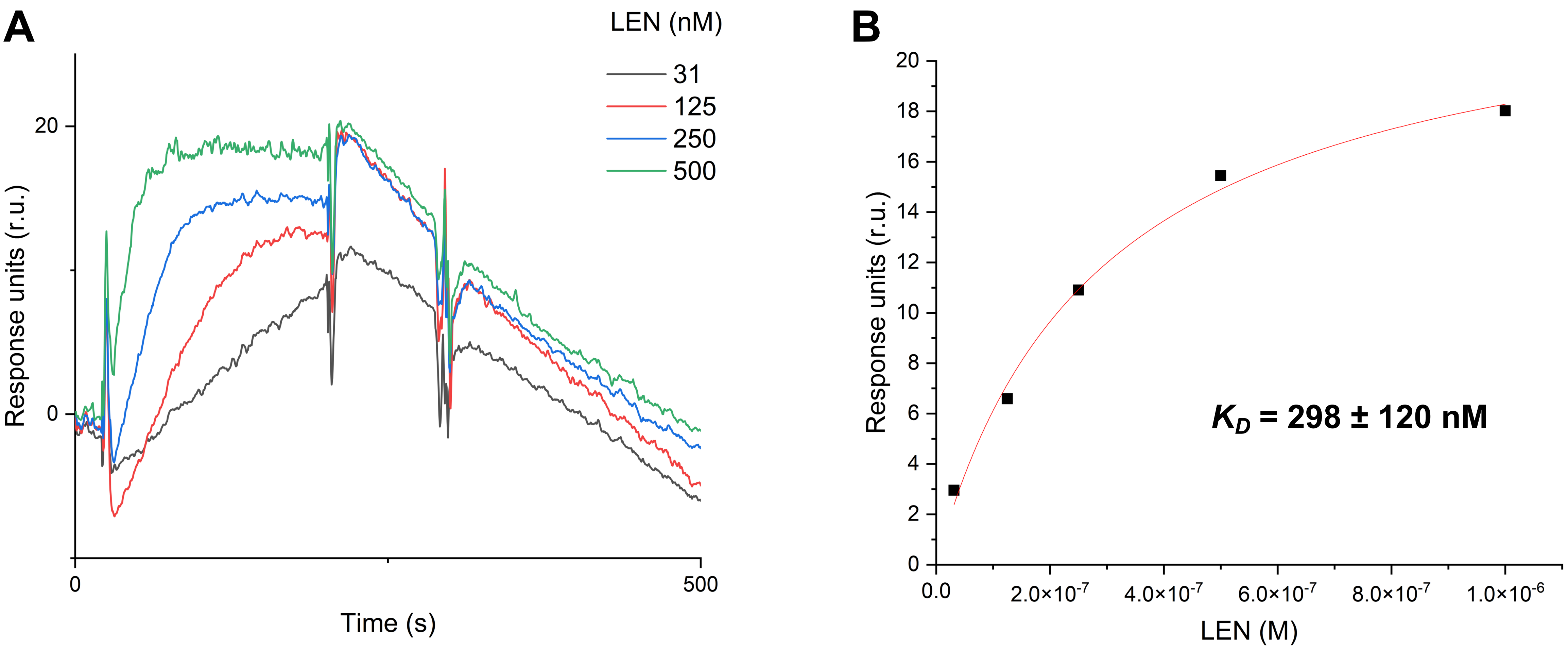


$k_{on}$ = 2.7 ± 1.1 x $10^4$ ($M^{-1}s^{-1}$)
$k_{off}$ = 9.5 ± 3.7 x $10^{-3}$ ($s^{-1}$)
$K_D$ = 361 ± 23 nM

**S4 Fig**

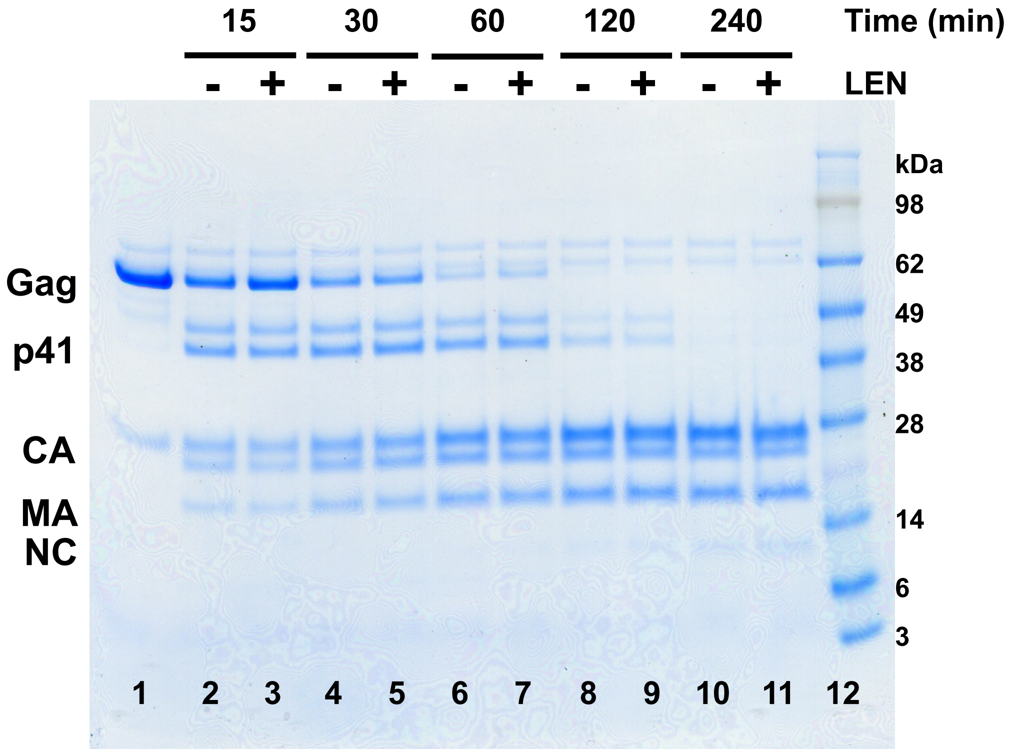
15
30
60
120
240
Time (min)
- + - + - + - + - +
LEN
kDa
98
62
49
38
28
14
6
3
Gag
p41
CA
MA
NC
1 2 3 4 5 6 7 8 9 10 11 12

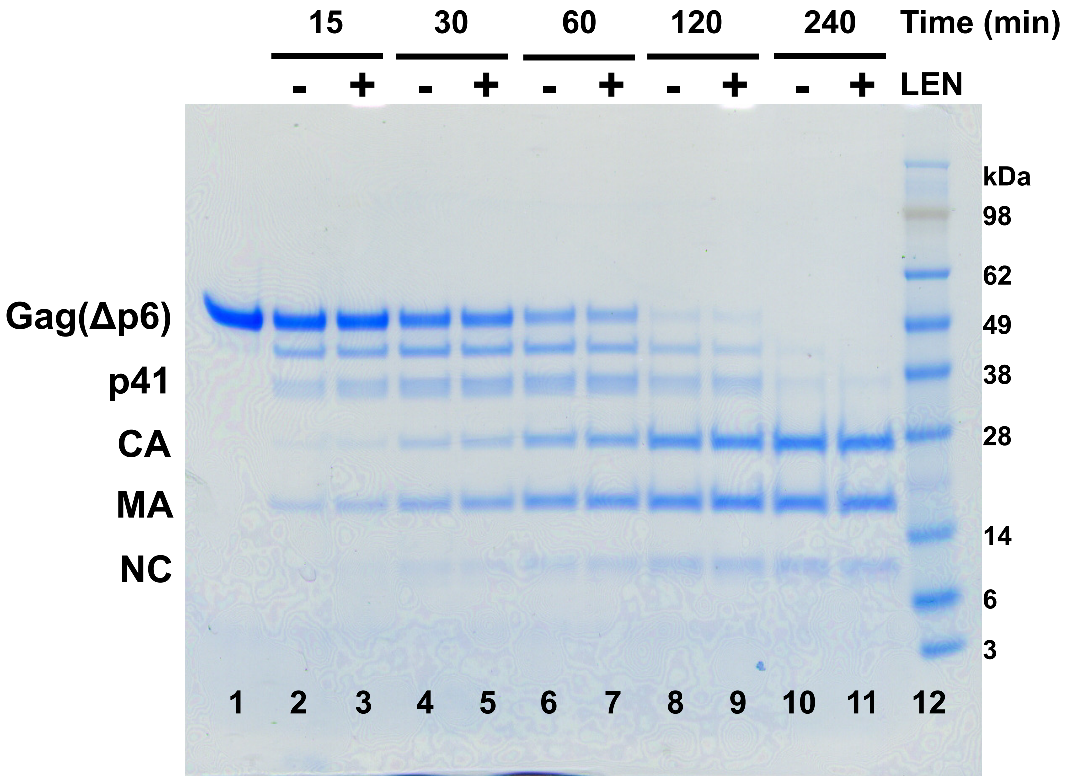
15
30
60
120
240
Time (min)
- + - + - + - + - +
LEN
kDa
98
62
49
38
28
14
6
3
Gag(Δp6)
p41
CA
MA
NC
1 2 3 4 5 6 7 8 9 10 11 12

S5 Fig

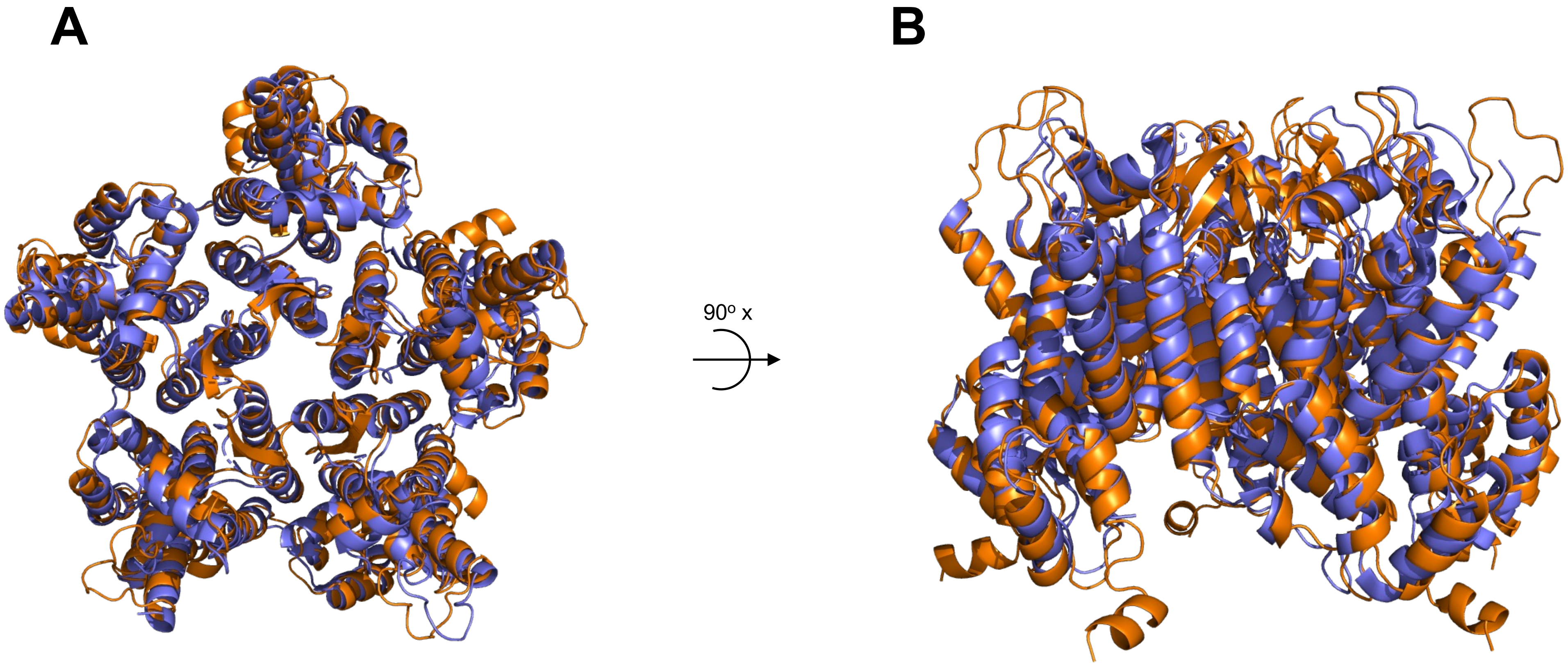


**RMSD: 1.04 Å over 208 residues**

**S6 Fig**

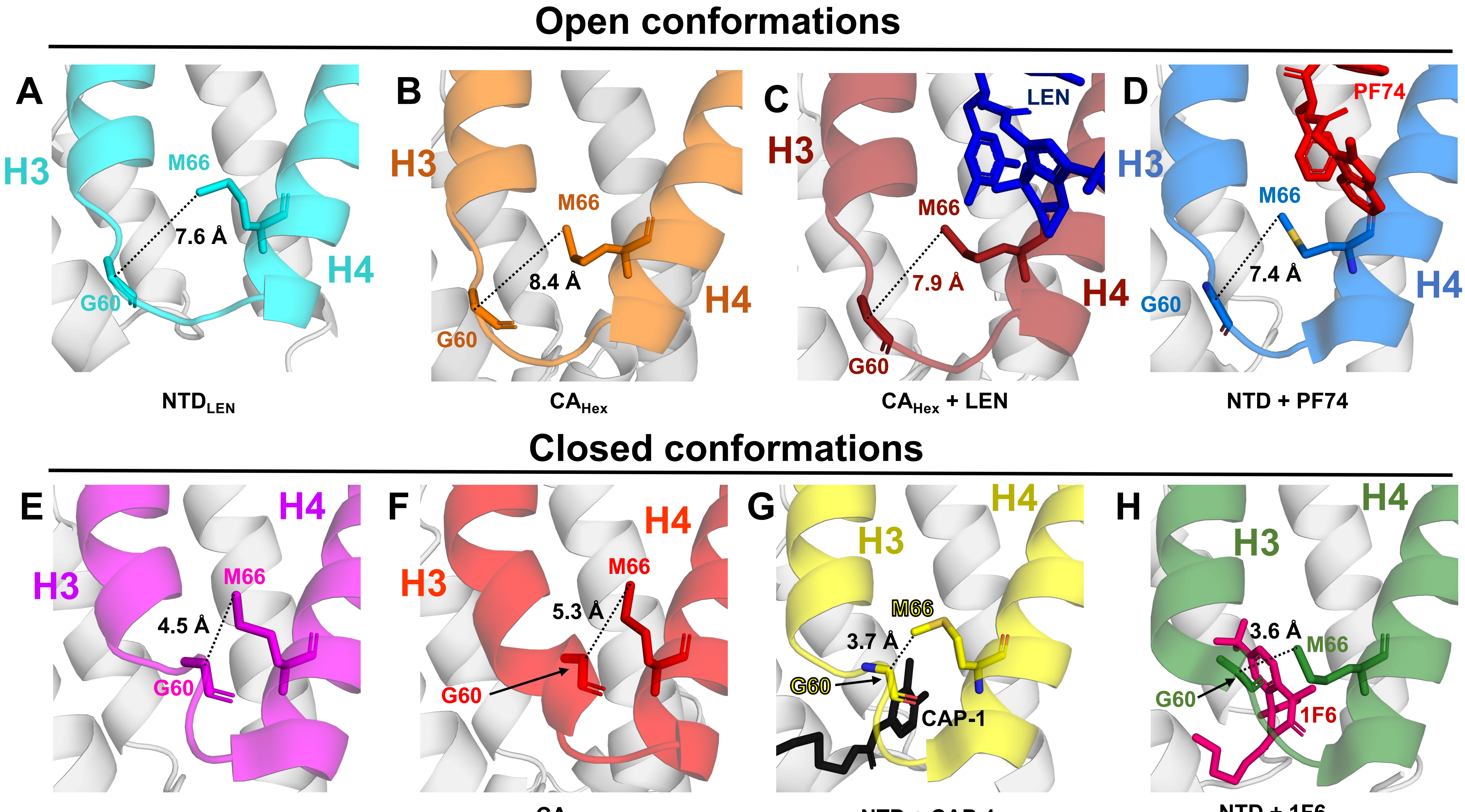
Open conformations
A
H3
M66
7.6 Å
H4
G60
NTD$_{LEN}$
B
H3
M66
8.4 Å
H4
G60
CA$_{Hex}$
C
LEN
H3
M66
7.9 Å
H4
G60
CA$_{Hex}$ + LEN
D
PF74
H3
M66
7.4 Å
H4
G60
NTD + PF74
Closed conformations
E
H4
H3
M66
4.5 Å
G60
NTD$_{Apo}$
F
H4
H3
M66
5.3 Å
G60
CA$_{Pnt}$
G
H4
H3
M66
3.7 Å
G60
CAP-1
NTD + CAP-1
H
H4
H3
3.6 Å
M66
G60
1F6
NTD + 1F6

S7 Fig

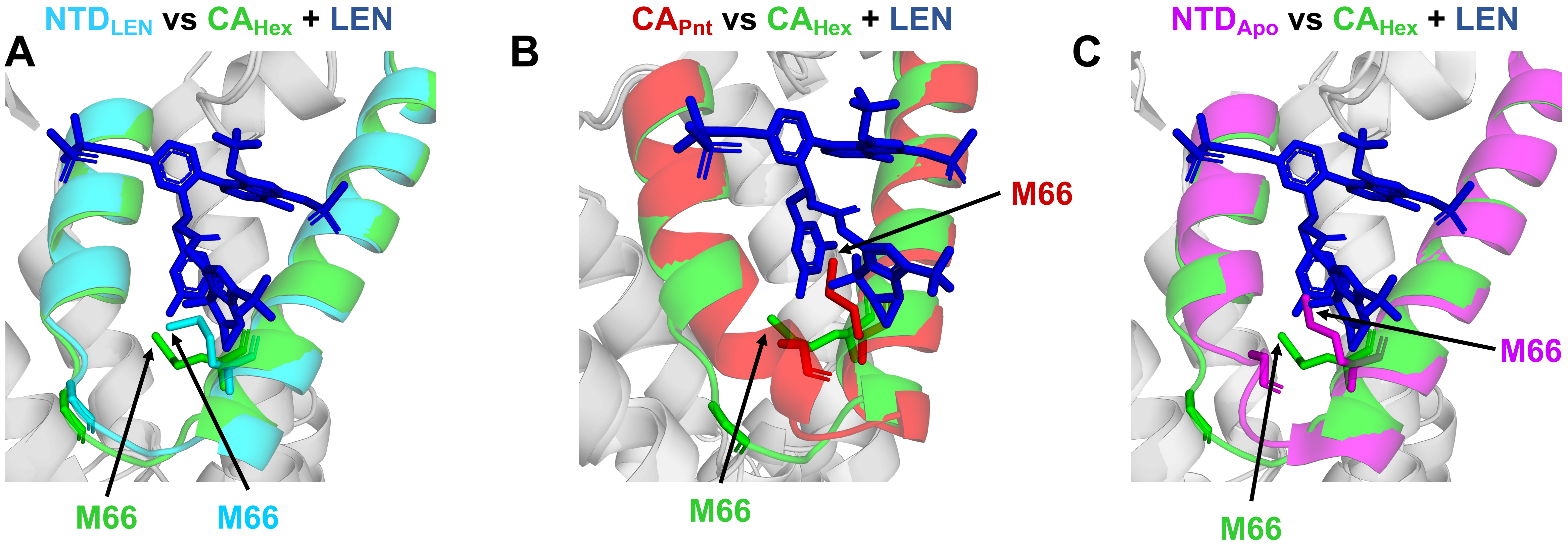
NTDLEN vs CAHex + LEN
A
M66
M66
CAPnt vs CAHex + LEN
B
M66
M66
NTDApo vs CAHex + LEN
C
M66
M66

S8 Fig
